# Resolving Three-State Ligand-Binding Mechanisms by Isothermal Titration Calorimetry: A Simulation Study

**DOI:** 10.1101/145516

**Authors:** Evgenii L. Kovrigin

## Abstract

In this paper, I theoretically analyzed ITC profiles for three-state equilibria involving ligand binding coupled to isomerization or dimerization transitions. Simulations demonstrate that the mechanisms where the free or ligand-bound protein undergoes dimerization (such that the ligand cannot bind to or dissociate from the dimer) produce very distinctive titration profiles. In contrast, profiles of the pre-existing equilibrium or induced-fit models cannot be distinguished from a simple two-state process, requiring data from additional techniques to positively identify these mechanisms.

---

Isothermal Titration Calorimetry (ITC) is a well-established technique for analysis of protein-ligand interactions to measure accurate binding affinity constants and establishing the reaction stoichiometry^*1*–*4*^. The ITC is frequently used to complement Nuclear Magnetic Resonance (NMR) studies to provide thermodynamic information on complex interaction mechanisms^*5*–*7*^. I previously reported analysis of NMR line shapes in titrations of systems with protein-ligand interactions coupled to isomerization or di-merization equlibria^*8*^. In this communication I am outlining major patterns one can expect in such three-state coupled equilibria from ITC experiments to help NMR and ITC experimentalists anticipate results from real systems.

## Methods

All simulations were performed using *LineShapeKin Simulation* software, which is designed to simulate NMR line shapes and ITC titration profiles. *LineShapeKin Simulation* was described in my previous report^*8*^ and is freely available from http://lineshapekin.net. The models under discussion in this paper are given in Fig. 1, where R stands for a “receptor”, which is a protein or nucleic acid or any other large or small molecule. The “L” stands for a ligand, which may as well be any of the above-mentioned molecules. We only need to discriminate R and L because in ITC one of the binding partners resides in the calorimeter cell, while another is added from the syringe. I am assigning “R” to the species in the cell and will casually refer to it as a “protein” while its binding partner, L, will be in a syringe and will be called a “ligand”. The reader must appreciate that this assignment is arbitrary and completely reversible. In fact, if one of the binding partners is poorly soluble it is advisable to place it in the calorimeter cell because the solution in the syringe has to be 10–20 times more concentrated. Such “reverse titrations” are described by the U-L and U-L2 models, which are equivalent to U-R and U-R2, therefore, will be dropped for clarity. Specific parameters for calculations of ITC profiles discussed in this paper are given in the Appendix.

**Fig. 1.**
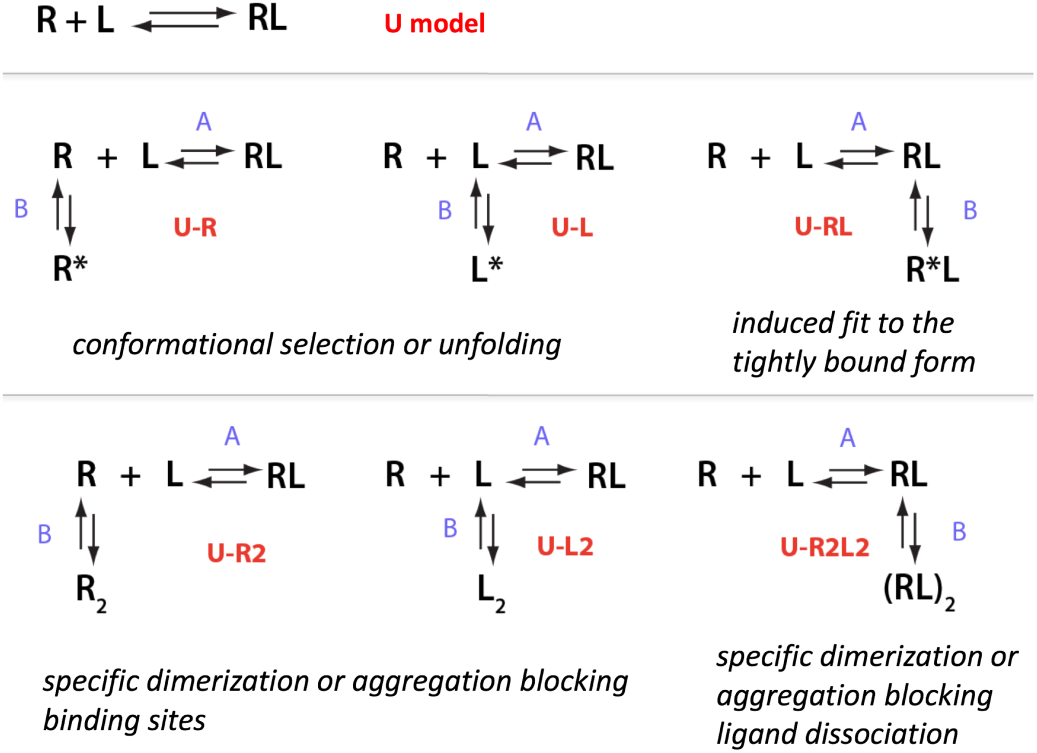
The three-state models evaluated in this study. Model names (according to a convention of the Li*neShapeKin Simulation*) are given in red. Binding and coupled transitions labels are blue.

## Results and Discussion

Fig. 1 summarizes the models that one can use to describe a protein molecule binding a single ligand and undergoing a one-step conformational or self-association transition. The real molecular events may be very complex involving multiple coupled transitions. The three-state models in Fig. 1 represent the first step towards this complexity; therefore, it is instructive to inspect titration patterns these systems may produce in ITC experiments (NMR line shapes for the same models have been described previously^*8*^).

The standard two-state “lock-and-key” binding mechanism of Emil Fischer is labeled as the U model in Fig. 1^*9*^. The U-R model is what is known as “pre-existing equilibrium”^*10*–*12*^, which may also be a folding-unfolding equilibrium coupled to ligand binding. In both cases, R is a binding-competent form, while R* is a conformation of the receptor that cannot bind L. Similarly, dimerization in U-R2 model produces a dimer that cannot bind a ligand. This is not a generic protein aggregation because the dimer formation *must block* the binding sites on both monomers. Likewise, the U-R2L2 model represents dimerization of the bound species such that the binding site is “locked” (buried or conformationally restrained), therefore, ligand cannot dissociate from the dimer. The dimer must dissociate first then the ligand may be released. The U-RL model is commonly known as Koshland’s induced-fit mechanism^*13*, *14*^. Thus, this set of three-state models approximates all major behaviors one can expect in molecular systems with one-to-one binding equilibria even if there are more steps ultimately involved in the process. For ITC analysis of molecules with multiple binding sites see an excellent treatment by Freiburger at al. ^*7*^.

In my simulations I assumed fast binding kinetics—millisecond time scale—faster than a typical time constant of the ITC instrument (on the order of 10 seconds). In this case, the ITC measurement is insensitive to kinetic parameters of the molecular mechanisms and only conveys thermodynamic information. Specific values for heat of formation of all species in the reaction mechanism were chosen to create situations when the heat effect of the coupled transition (B transition) *subtracts* from or *adds* to the heat of the ligand binding process.

Fig. 2 demonstrates the effect of coupled transitions on the ITC profiles. A simple 1:1 binding mechanism (U model) with K_a_=10^6^ M^−1^ serves as a reference (solid curves). The sample concentration in the ITC cell is chosen based on a value of c=K_a_*[in cell]*n, where K_a_ is the equilibrium association constant, and n is the stoichiometry of a binding reaction. An optimal range of c is between 1 and 1000^*1*, *15*^, therefore c-value was set 100 by choosing a total concentration of R in the cell equal to 0.1 mM. Equilibrium constants in the B transition were chosen to create a ratio of isomers or monomers/dimers (in absence of L for U-R2, and under saturating concentrations of L for U-R2L2) close to unity.

**Fig. 2.**
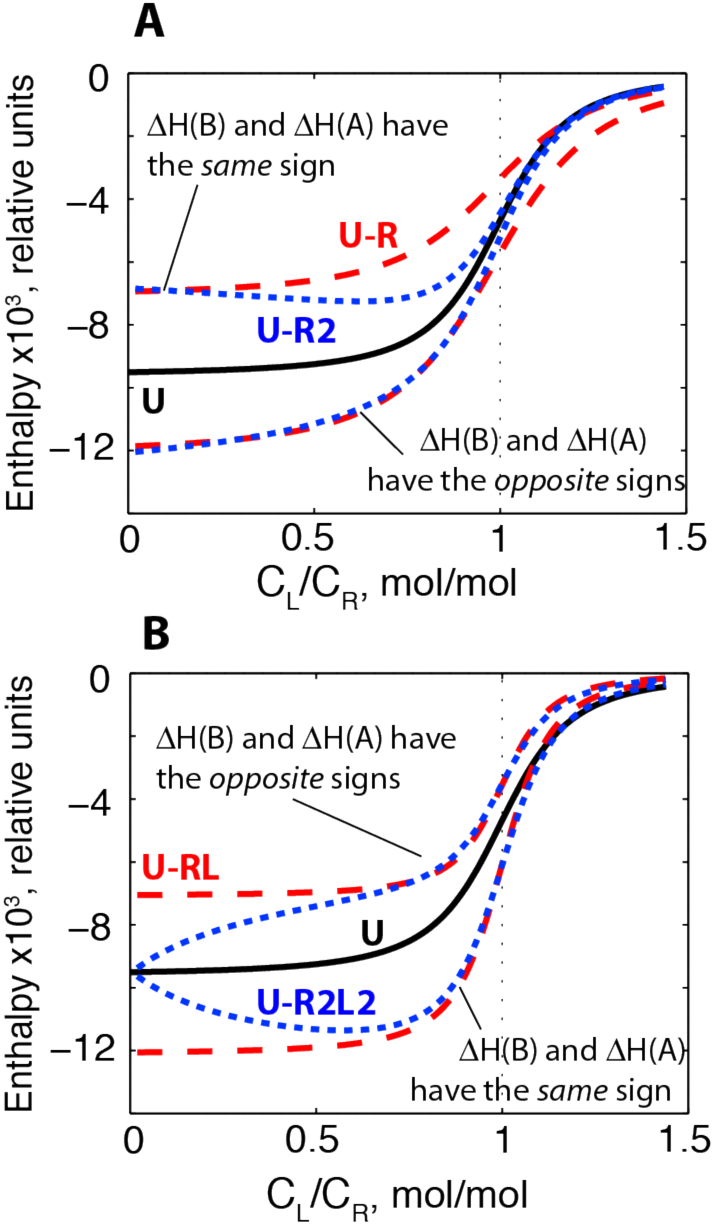
Effect of isomerization and dimerization equilibria on ITC titration profiles in three-state systems. Black solid curves in both panels correspond to the U model with an association constant of 10^6^ M^−1^ and the binding enthalpy of −1 kJ/mol (chosen as a reference value). Total concentration of R in all cases is 0.1 mM. (**Panel A**) Profiles for the U-R model (red dashed) and U-R2 model (blue dotted) plotted with the enthalpy of a B transition set to −0.5 kJ/mole (U-R and U-R2 curves above the U-model curve) and 0.5 kJ/mole (below). (**Panel B**) Profiles for the U-RL model (red dashed) and U-R2L2 model (blue dotted) plotted with the enthalpy of a B transition set to −0.5 kJ/mole (U-RL and U-R2L2 curves below the U-model curve) and 0.5 kJ/mole (above).

The simulations for pre-existing equilibria U-R and U-R2 demonstrated a couple of distinctive trends (Fig. 2.A, dashed and dotted curves). In the first place, we observe that the B-transition heat is *subtracted* from the ligand binding heat (transition A) if ΔH(A) and ΔH(B) have the *same sign* (curves lie above the solid U-model curve). This is a natural consequence of the fact that R* and R_2_ are non-binding species; therefore, upon ligand addition the process of formation of R* or R_2_ is reversed. Conversely, when ΔH(A) and ΔH(B) have *opposite* signs the overall heat effect is increased (curves lie below the U-model curve).

The second notable consequence of the R* ⇔ R equilibrium coupled to binding is that formation of R* species *opposes* formation of the complex resulting in a more shallow sigmoidal shape for U-R. The end result is that U-R model profiles (dashed curves) appear as scaled-up or down versions of the U-model curve (solid) with a reduced apparent equilibrium binding constant. The U-R2 profiles are more complex because of the concentration dependence of R_2_ formation. During the titration, the amount of free R is gradually depleted and R_2_ ⇔ 2R equilibrium progressively shifts towards R (while R* ⇔ R equilibrium is *unaffected* by a change in concentration of R so R/R* ratio remains constant at all times). The U-R2 model (dotted curves) behaves like U-R in the beginning of a titration when the initial high concentration of free R allows significant amounts of R_2_ to be formed. As C_L_/C_R_ increases the U-R2 profile gradually approaches the U-model profile in the near-equivalence region where R is sufficiently depleted to form negligible amounts of R_2_. One may expect that a simultaneous fitting of data from two ITC experiments performed with different total concentration of receptor in the calorimeter cell would be sufficient to completely discriminate between the U-R and U-R2 mechanisms [provided ΔH(B) is comparable to ΔH(A)].

The models with isomerization or dimerization occurring *after* formation of the RL complex are analyzed in the Fig. 2, panel B. Formation of (RL)* or (RL)_2_ species affects the total binding heat in a fashion similar to the pre-existing equilibria in R with the only difference that here ΔH(B) adds to ΔH(A) if they have the same signs (curves lie below the U model) and subtracts if the signs are opposite (curves above the U-model). Similarly to the U-R case, the isomerization equilibrium in U-RL model does not depend on RL concentration and, therefore, the titration profiles strongly resemble the U-model curve with effective affinity increased by the coupled isomerization equilibrium. The U-R2L2 curves reflect concentration dependence of the (RL)_2_ formation: very little dimer is formed in the beginning when RL concentration is still small; formation of (RL)_2_ is more and more favored upon titration progress. As a result, the U-R2L2 curves begin from the U-model curve, then deviate and, eventually, approach the U-RL curves when the dimer-monomer ratio reaches the R*/R ratio of the U-RL simulation.

In summary, we observe that the ligand binding to proteins coupled to *isomerization* equilibria in either ligand-free or ligand-bound states cannot produce readily identifiable patterns in the ITC measurements. Therefore, pre-existing equilibrium (or folding/unfolding) and induced fit titration profiles cannot be differentiated from a simple 1:1 binding event by ITC. In other words, the three-state ITC curve can be perfectly fit to the standard two-state binding curve producing effective enthalpies and binding constants while giving no hint of a more complex equilibrium taking place. In contrast, binding of ligands coupled to *dimerization* (and, correspondingly—oligomerization) of the free and bound receptor produces titration profiles remarkably different from the simple two-state model. We may conclude that U-R2 and U-R2L2 models should be discernible by comparative fitting of experimental data to two- and three-state models.

## ACKNOWLEDGMENT

The author is indebted to Dr. James Kempf for helpful discussions of this material. The author acknowledges financial support of the Marquette University Committee on Research (2012 Summer Faculty Fellowship and 2012 Regular Research Grant).

## Appendix: Simulation Parameters

The following are the parameters of *LineShapeKin Simulation* that one can use to reproduce ITC profiles in given in Figure 2. For installation instructions see http://lineshapekin.net/-LineShapeKinSimulation411. To run a simulation, save setup parameters in “setup.txt”, desired model parameters—in “model_parameters.txt” files, and type on the MATLAB command line: “Simulate setup model_parameters”. A technical note: the *LineShapeKin Simulation* requires setting *absolute* enthalpies for individual species to compute the ΔH of transitions A and B (as contrasted to directly setting binding and isomerization/dimerization transition enthalpy change, ΔH). By historical reasons, the absolute values are denoted by “dH” in the parameter files.

### Setup file

~~~
# LineShapeKin Simulation
#
# General simulation settings

# ======= Folder to put simulation results into ==========
Results_Path Simulation_Results

# ======== Total Ligand to total Receptor molar ratio at each titration point ========
# NOTE: first point L/R=0 has to be non-zero very small number to prevent /0

LRratio   1e-3 0.1 0.2 0.3 0.4 0.5 0.6 0.7 0.8 0.9  1.0  1.1 1.2

# ======== Number of points on smooth population curves and on ITC profile ========
# NOTE: >100 points will make complex models slow!

LRpoints 150

# ======== constant M_ligand/Receptor ratio ========
# (in models that use it, otherwise - ignored)

MRratio  0

# ======== Concentration of receptor, M ========
# You may have Rtotal vary corresponding to your experimental conditions and all calculations
# at specific titration points given by LRratio will be correct. However,
# the smooth curves will only be computed for fixed Rtotal.

Rtotal   1e-4  1e-4  1e-4  1e-4  1e-4  1e-4  1e-4  1e-4  1e-4  1e-4  1e-4  1e-4  1e-4

# ========== Plotting options ==============

# – number of points in each individual spectral trace
Spectral_Points   300

#– limits of a spectral window /s
w_min   -50
w_max   350

#— YES to display the titration series stacked rather than overlayed
Stacked_Plot  YES

#– if Stacked_Plot=YES, this is the percentage of the peak height for shifting the next trace up
%
Percent_Shift  15
~~~

### Model file for lock-and-key mechanism

~~~
# Unique model identifier
Model_code     U

# Model description
Description  Simple  1:1

# Association constants
Ka_names  A
Ka        1e6

# Rate constants of REVERSE reactions
k_names    A
k2         500

# Names of NMR-active species
Species_names     R   RL

# Names of NMR unobservable species
NMR_invisible_species_names    L

# Chemical shifts of pure species, 1/s
w0        0       300

# Relaxation rates of pure species, 1/s
R2      10      10

# Heat of formation of the species, relative units
# The original species is a standard state with dH=0 !
dH     0   -1
~~~

### Model file for preexisting equlibrium mechanism with negative enthalpy of B transition

~~~
# Unique model identifier
Model_code    U_R

# Model description
Description  Fast binding, fast isomerization

# Association constants
Ka_names   A      B
Ka         1e6    1.1

# Rate constants of REVERSE reactions
k_names     A   B
k2         500  500

# Names of NMR-active species
Species_names      R   R*   RL

# Names of NMR unobservable species
NMR_invisible_species_names    L

# Chemical shifts of pure species, 1/s
w0        150   0    300

# Relaxation rates of pure species, 1/s
R2      10      10     10

# Heat of formation of the species, relative units
# The original species is a standard state with dH=0 !
dH     0    -0.5    -1
~~~

### Model file for preexisting equlibrium mechanism with positive enthalpy of B transition

~~~
# Unique model identifier
Model_code     U_R

# Model description
Description  Fast binding, fast isomerization

# Association constants
Ka_names   A      B
Ka         1e6    1.1

# Rate constants of REVERSE reactions
k_names     A    B
k2         500   500

# Names of NMR-active species
Species_names      R   R*   RL

# Names of NMR unobservable species
NMR_invisible_species_names     L

# Chemical shifts of pure species, 1/s
w0      150  0    300

# Relaxation rates of pure species, 1/s
R2      10      10     10

# Heat of formation of the species, relative units
# The original species is a standard state with dH=0 !
dH     0    0.5    -1
~~~

### Model with dimerization of free protein with negative enthalpy of B transition

~~~
# Unique model identifier
Model_code     U_R2

# Model description
Description  Fast binding, fast dimerization

# Association constants
Ka_names   A      B
Ka         1e6    5e3

# Rate constants of REVERSE reactions
k_names     A    B
k2          500    500

# Names of NMR-active species
Species_names      R   RR    RL

# Names of NMR unobservable species NMR_invisible_species_names     L

# Chemical shifts of pure species, 1/s
w0         150   0    300

# Relaxation rates of pure species, 1/s
R2      10      10     10

# Heat of formation of the species, relative units
# The original species is a standard state with dH=0 !
dH     0     -0.5   -1
~~~

### Model with dimerization of free protein with positive enthalpy of B transition

~~~
# Unique model identifier
Model_code     U_R2

# Model description
Description  Fast binding, fast dimerization

# Association constants
Ka_names   A      B
Ka         1e6    5e3

# Rate constants of REVERSE reactions
k_names    A     B
k2         500     500

# Names of NMR-active species
Species_names      R   RR   RL

# Names of NMR unobservable species
NMR_invisible_species_names     L

# Chemical shifts of pure species, 1/s
w0         150   0   300

# Relaxation rates of pure species, 1/s
R2      10      10     10

# Heat of formation of the species, relative units
# The original species is a standard state with dH=0 !
dH     0    0.5    -1
~~~

### Model file for induced mechanism with negative enthalpy of B transition

~~~
# Unique model identifier
Model_code     U_RL

# Model description
Description  Fast binding, fast isomerization

# Association constants
Ka_names   A      B
Ka         1e6    1.1

# Rate constants of REVERSE reactions
k_names     A     B
k2         500   500

# Names of NMR-active species
Species_names      R   RL   R*L

# Names of NMR unobservable species
NMR_invisible_species_names     L

# Chemical shifts of pure species, 1/s
w0        0   150    300

# Relaxation rates of pure species, 1/s
R2      10      10     10

# Heat of formation of the species, relative units
# The original species is a standard state with dH=0 !
dH     0    -1   -1.5
~~~

### Model file for induced mechanism with positive enthalpy of B transition

~~~
# Unique model identifier
Model_code     U_RL

# Model description
Description  Fast binding, fast isomerization

# Association constants
Ka_names   A      B
Ka         1e6    1.1

# Rate constants of REVERSE reactions
k_names     A     B
k2         500  500

# Names of NMR-active species
Species_names      R   RL   R*L

# Names of NMR unobservable species
NMR_invisible_species_names     L

# Chemical shifts of pure species, 1/s
w0        0   150    300

# Relaxation rates of pure species, 1/s
R2      10      10    10

# Heat of formation of the species, relative units
# The original species is a standard state with dH=0 !
dH     0    -1    -0.5
~~~

### Model file for dimerization of the bound state with negative enthalpy of B transition

~~~
# Unique model identifier
Model_code     U_R2L2

# Model description
Description Fast binding, fast dimerization

# Association constants
Ka_names   A      B
Ka         1e6    5e3

# Rate constants of REVERSE reactions
k_names     A     B
k2          500     500

# Names of NMR-active species
Species_names      R   RL   R2L2

# Names of NMR unobservable species
NMR_invisible_species_names    L

# Chemical shifts of pure species, 1/s
w0         0   150   300

# Relaxation rates of pure species, 1/s
R2      10      10     10

# Heat of formation of the species, relative units
# The original species is a standard state with dH=0 !
dH     0    -1    -1.5
~~~

### Model file for dimerization of the bound state with positive enthalpy of B transition

~~~
# Unique model identifier
Model_code     U_R2L2

# Model description
Description Fast binding, fast dimerization

# Association constants
Ka_names   A      B
Ka         1e6    5e3

# Rate constants of REVERSE reactions
k_names     A    B
k2          500    500

# Names of NMR-active species
Species_names      R   RL   R2L2

# Names of NMR unobservable species
NMR_invisible_species_names    L

# Chemical shifts of pure species, 1/s
w0         0   150    300

# Relaxation rates of pure species, 1/s
R2      10      10     10

# Heat of formation of the species, relative units
# The original species is a standard state with dH=0 !
dH     0    -1    -0.5
~~~

## REFERENCES

[1] Ladbury, J. E., and Chowdhry, B. Z. (1996) Sensing the heat: the application of isothermal titration calorimetry to thermodynamic studies of biomolecular interactions, Chem Biol 3, 791–801.

[2] Velazquez-Campoy, A., Ohtaka, H., Nezami, A., Muzammil, S., and Freire, E. (2004) Isothermal titration calorimetry, Curr Protoc Cell Biol Chapter 17, Unit 17 18.

[3] Freyer, M. W., and Lewis, E. A. (2008) Isothermal titration calorimetry: experimental design, data analysis, and probing macromolecule/ligand binding and kinetic interactions, Methods Cell Biol 84, 79–113.

[4] Ward, W. H., and Holdgate, G. A. (2001) Isothermal titration calorimetry in drug discovery, Prog Med Chem 38, 309–376.

[5] Korzhnev, D. M., Bezsonova, I., Lee, S., Chalikian, T. V., and Kay, L. E. (2009) Alternate binding modes for a ubiquitin-SH3 domain interaction studied by NMR spectroscopy, J Mol Biol 386, 391–405.

[6] Demers, J. P., and Mittermaier, A. (2009) Binding mechanism of an SH3 domain studied by NMR and ITC, J Am Chem Soc 131, 4355–4367.

[7] Freiburger, L., Auclair, K., and Mittermaier, A. (2015) Global ITC fitting methods in studies of protein allostery, Methods 76, 149–161.

[8] Kovrigin, E. L. (2012) NMR line shapes and multi-state binding equilibria, J Biomol NMR 53, 257–270.

[9] Fischer, E. (1894) Einfluss der Configuration auf die Wirkung der Enzyme (Influence of Configuration on the Effect of Enzymes), Ber. Dtsch. Chem. Ges. 27, 2984–2993.

[10] Monod, J., Wyman, J., and Changeux, J. P. (1965) On the Nature of Allosteric Transitions: a Plausible Model., J Mol Biol. 12, 88–118.

[11] Kumar, S., Ma, B. Y., Tsai, C. J., Sinha, N., and Nussinov, R. (2000) Folding and binding cascades: Dynamic landscapes and population shifts, Protein Science 9, 10–19.

[12] Gunasekaran, K., Ma, B., and Nussinov, R. (2004) Is allostery an intrinsic property of all dynamic proteins?, Proteins 57, 433 – 443.

[13] Koshland, D. E. (1958) Application of a theory of enzyme specificity to protein synthesis, Proc Natl Acad Sci U S A 44, 98–104.

[14] Koshland, D. E. (1994) The key-lock theory and the induced fit theory, Angewandte Chemie International Edition Engl. 33, 2375–2378.

[15] Wiseman, T., Williston, S., Brandts, J. F., and Lin, L. N. (1989) Rapid Measurement of Binding Constants and Heats of Binding Using a New Titration Calorimeter, Analytical Biochemistry 179, 131–137.

